# Chronic stroke sensorimotor impairment is related to smaller hippocampal volumes: An ENIGMA analysis

**DOI:** 10.1101/2021.10.26.465924

**Authors:** A Zavaliangos-Petropulu, B Lo, MR Donnelly, N Schweighofer, Keith Lohse, Neda Jahanshad, G Barisano, N Banaj, MR Borich, LA Boyd, CM Buetefisch, WD Byblow, JM Cassidy, CC Charalambous, AB Conforto, JA DiCarlo, AN Dula, N Egorova-Brumley, MR Etherton, W Feng, KA Fercho, F Geranmayeh, CA Hanlon, KS Hayward, B Hordacre, SA Kautz, MS Khlif, H Kim, A Kuceyeski, DJ Lin, M Lotze, J Liu, BJ MacIntosh, JL Margetis, F Piras, A Ramos-Murguialday, KP Revill, PS Roberts, AD Robertson, HM Schambra, NJ Seo, MS Shiroishi, SR Soekadar, G Spalletta, M Taga, WK Tang, GT Thielman, D Vecchio, NS Ward, LT Westlye, E Werden, C Winstein, GF Wittenberg, SL Wolf, KA Wong, C Yu, A Brodtmann, SC Cramer, PM Thompson, S-L Liew

**Affiliations:** Mark and Mary Stevens Neuroimaging and Informatics Institute, Keck School of Medicine, University of Southern California, Los Angeles, CA, 91001, USA; Neuroscience Graduate Program, University of Southern California, Los Angeles, CA, 90089, USA; Chan Division of Occupational Science and Occupational Therapy, University of Southern California, Los Angeles, CA, 90089, USA; Biokinesiology and Physical Therapy, University of Southern California, Los Angeles, CA, 90089, USA; Physical Therapy and Neurology, Washington University School of Medicine in Saint Louis, St. Louis, Missouri, 63110, USA; Laboratory of Neuropsychiatry, IRCCS Santa Lucia Foundation, Rome, 00167, Italy; Department of Rehabilitation Medicine, Division of Physical Therapy, Emory University School of Medicine, Atlanta, 30322, USA; Department of Physical Therapy, University of British Columbia, Vancouver, V6T1Z3, Canada; Department of Neurology, SOM, Emory University, Atlanta, Georgia, 30322, USA; Department of Exercise Sciences, and Centre for Brain Research, University of Auckland, Auckland, 92019, New Zealand; Department of Allied Health Sciences, University of North Carolina at Chapel Hill, Chapel Hill, North Carolina, 27516, USA; Department of Basic and Clinical Sciences, University of Nicosia Medical School, Nicosia, 2414, Cyprus; Center for Neuroscience and Integrative Brain Research (CENIBRE), Nicosia, 2414, Cyprus; Hospital das Clínicas, São Paulo University, São Paulo, 05403000, Brazil; Hospital Israelita Albert Einstein, São Paulo, 01431000, Brazil; Center for Neurotechnology and Neurorecovery (CNTR), Massachusetts General Hospital, Boston, 02114, Massachusetts, USA; Department of Neurology, Dell Medical School, University of Texas at Austin, Austin, Texas, 78712, USA; Melbourne School of Psychological Sciences, University of Melbourne, Parkville, Victoria, 3052, Australia; J. Philip Kistler Stroke Research Center, Department of Neurology, Massachusetts General Hospital, Boston, Massachusetts, 02130, USA; Department of Neurology, Duke University School of Medicine, Durham, North Carolina, 27710, USA; Basic Biomedical Sciences, University of South Dakota, Vermillion, South Dakota, 57069, USA; Federal Aviation Administration, Civil Aerospace Medical Institute, Oklahoma City, OK, 73125, USA; Brain Sciences, Imperial College London, London, W12 0NN, United Kingdom; Wake Forest School of Medicine, Winston Salem, North Carolina, 27157, USA; Departments of Physiotherapy and Medicine, Heidelberg, Victoria, 3084, Australia; The Florey Institute of Neuroscience and Mental Health, Heidelberg, Victoria, 3084, Australia; Innovation, Implementation and Clinical Translation (IIMPACT) in Health, Allied Health and Human Performance, University of South Australia, Adelaide, South Australia, 5000, Australia; Ralph H Johnson Veterans Affairs Medical Center, Charleston, South Carolina, 29401, USA; Department of Health Sciences & Research, Medical University of South Carolina, Charleston, South Carolina, 29401, USA; Department of Radiology, Weill Cornell Medicine, New York, New York, 10065, USA; Functional Imaging, Institute for Diagnostic Radiology and Neuroradiology, University Medicine Greifswald, 17475 Germany; Department of Radiology, Tianjin Medical University General Hospital, Tianjin, China; Hurvitz Brain Sciences Program, Sunnybrook Research Institute, Toronto, M4N3M5, Canada; Department of Medical Biophysics, University of Toronto, Toronto, Ontario, M5S1A1,Canada; University of Southern California, Los Angeles, CA, 90089, USA; Institute of Medical Psychology and Behavioral Neurobiology, University of Tübingen, Tübingen, 72076, Germany; Health Division, TECNALIA, San Sebastian, 20009, Spain; Department of Psychology, Emory University, Atlanta, Georgia, 30322, USA; Department of Physical Medicine and Rehabilitation, Cedars-Sinai, Los Angeles, CA, 90048, USA; Department of Kinesiology and Health Sciences, University of Waterloo, Waterloo, Ontario N2L3G1, Canada; Departments of Neurology & Rehabilitation Medicine, NYU Langone, New York, New York, 10017, USA; Department of Rehabilitation Sciences, Department of Health Science and Research, Medical University of South Carolina, Charleston, South Carolina, 29425, USA; Department of Radiology, Keck School of Medicine, University of Southern California, Los Angeles, CA, 91001, USA; Clinical Neurotechnology Laboratory, Dept. of Psychiatry and Neurosciences (CCM), Charité - Universitätsmedizin Berlin, Berlin, 10117, Germany; NYU Langone Department of Neurology, New York, New York, 10017, USA; Department of Psychiatry, the Chinese University of Hong Kong, Hong Kong, China; University of the Sciences Department of Physical Therapy and Neuroscience, Philadelphia, Pennsylvania, 19104, USA; University College London Queen Square Institute of Neurology, London, SE59AX, United Kingdom; Department of Psychology, University of Oslo, Olso, 0317, Norway; Department of Mental Health and Addiction, Oslo University Hospital, Oslo, 0455, Norway; Melbourne Dementia Research Center, University of Melbourne, Victoria, 3052, Australia; Department of Neurology, University of Pittsburgh, Pittsburgh, Pennsylvania 15213, USA; Department of Veterans Affairs, Geriatrics Research Educational & Clinical Center, VAPHS, Pittsburgh, Pennsylvania, 15240, USA; Department of Medicine, Emory University School of Medicine, Atlanta, Georgia, 30322, USA; Department of Physical Medicine & Rehabilitation, Dell Medical School, University of Texas at Austin, Austin, Texas, 78701, USA; Department of Neurology, University of California Los Angeles, David Geffen School of Medicine, Los Angeles, CA, 90095, USA; California Rehabilitation Hospital, Los Angeles, CA, 90067, USA

**Keywords:** Stroke, Sensorimotor Impairment, Hippocampus, MRI

## Abstract

Persistent sensorimotor impairments after stroke can negatively impact quality of life. The hippocampus is involved in sensorimotor behavior but has not been widely studied within the context of post-stroke upper limb sensorimotor impairment. The hippocampus is vulnerable to secondary degeneration after stroke, and damage to this region could further weaken sensorimotor circuits, leading to greater chronic sensorimotor impairment. The purpose of this study was to investigate the cross-sectional association between non-lesioned hippocampal volume and upper limb sensorimotor impairment in people with chronic stroke. We hypothesized that smaller ipsilesional hippocampal volumes would be associated with worse upper-limb sensorimotor impairment.

Cross-sectional T1-weighted brain MRIs were pooled from 357 participants at the chronic stage after stroke (>180 days post-stroke) compiled from 18 research cohorts worldwide in the ENIGMA Stroke Recovery Working Group (age: median = 61 years, interquartile range = 18, range = 23-93; 135 women and 222 men). Sensorimotor impairment was estimated from the Fugl-Meyer Assessment of Upper Extremity scores. Robust mixed-effects linear models were used to test associations between post-stroke sensorimotor impairment and hippocampal volumes (ipsilesional and contralesional separately; Bonferroni-corrected, *p*-*value* < 0.025), controlling for age, sex, lesion volume, and lesioned hemisphere. We also performed an exploratory analysis to test whether sex differences influence the relationship between sensorimotor impairment and hippocampal volume.

Upper limb sensorimotor impairment was positively associated with ipsilesional (*p* = 0.005; *d* = 0.33) but not contralesional (*p* = 0.96; *d* = 0.01) hippocampal volume, such that impairment was worse for participants with smaller ipsilesional hippocampal volume. This association remained significant independent of lesion volume or other covariates (*p* = 0.001; *d* = 0.36). Evidence indicates an interaction between sensorimotor impairment and sex for both ipsilesional (*p* = 0.008; *d* = −0.29) and contralesional (*p* = 0.006; *d* = −0.30) hippocampal volumes, whereby women showed progressively worsening sensorimotor impairment with smaller hippocampal volumes compared to men.

The present study has identified a novel association between chronic post-stroke sensorimotor impairment and ipsilesional, but not contralesional, hippocampal volume. This finding was not due to lesion size and may be stronger in women. We also provide supporting evidence that smaller hippocampal volume post-stroke is likely a consequence of ipsilesional damage, which could provide a link between vascular disease and other disorders, such as dementia.

## Introduction

Sensorimotor impairments are a major burden of disease for stroke survivors^1^. To help clinicians, caregivers, and patients make informed and effective rehabilitation treatment decisions, there is a critical need to identify biomarkers that accurately predict a patient’s potential for sensorimotor recovery^2,3^. MRI studies of regional brain volumes suggest secondary degeneration of adjacent or remote regions may contribute to sensorimotor impairment and could influence post-stroke sensorimotor outcomes^4,5^.

The hippocampus is a brain region that is particularly vulnerable to post-stroke secondary degeneration. Both rodent^6^ and human^7–11^ stroke studies have shown evidence of damage within the hippocampus in the same hemisphere as an infarct (ipsilesional), but outside of the lesion itself. Using structural MRI, smaller ipsilesional hippocampal volumes in stroke patients have been reported in comparison to healthy controls^7–11^, as well as smaller bilateral hippocampal volumes at the time of stroke^12^ and accelerated hippocampal atrophy observed most prominently three months after stroke^6,11^. Studies have also reported magnetic resonance spectroscopy evidence of contralesional hippocampal neuronal loss^10^ and contralesional hippocampal atrophy measured with longitudinal MRI^11^. Given that stroke-related infarctions of the hippocampus are uncommon^13,14^, post-stroke hippocampal atrophy is most likely attributed to secondary degenerative mechanisms such as spreading depression^6^ or reduced connectivity to lesioned structures^13^, among others. However, the extent to which lesion volume relates to hippocampal damage remains unclear.

The hippocampus is widely known for its key role in memory, and this has led the field of stroke recovery research to primarily focus on the role of hippocampal damage in cognitive impairment^6,7,10^. Although not typically considered a primary sensorimotor region, there is evidence that the hippocampus is also involved in sensorimotor behavior. The hippocampus is densely connected to brain areas that play an important role in sensorimotor processing such as the thalamus and basal ganglia through the spinal-limbic pathway^15^. Reports of hippocampal activity during sensorimotor behavior such as sensorimotor integration^16^, sensorimotor learning^17–19^, and motor control^18^ suggest that the hippocampus plays a role in sensorimotor circuits. Sensorimotor task-related functional connectivity with the hippocampus has also been reported with the thalamus^20^, sensorimotor cortex^18^, and the supplementary motor area^21^. However, the relationship between hippocampal structural integrity and post-stroke upper-limb sensorimotor impairment remains unclear. Given the involvement of the hippocampus in sensorimotor circuits, hippocampal damage due to secondary degeneration after stroke could further weaken sensorimotor circuits, leading to greater chronic sensorimotor impairment. Alternatively, damage to the thalamus, basal ganglia, sensorimotor cortex, or supplementary motor area, which are typically associated with greater sensorimotor impairment, may lead to downstream degeneration of the hippocampus through functional or structural connections.

Dementia studies^22^ and healthy ageing^23^ research also suggest that hippocampal atrophy may differ by sex, as hippocampal atrophy has been found to accelerate in post-menopausal women. Estrogen levels may play a mediating role in these trends^24^ and have been associated with stroke severity and mortality^25^. Stroke-related outcomes including disability and quality of life are generally poorer in women than men^1,26,27^, although conclusive sex differences have not been reported in terms of post-stroke sensorimotor impairment^28^. As such, sex could moderate the relationship between sensorimotor impairment and hippocampal volume following a stroke. In particular, women may have smaller hippocampal volumes and worse sensorimotor impairment, potentially leading to stronger effect sizes compared to men.

In addition, associations between lesion size and hippocampal volume remain unclear. One study reports larger lesion size is directly associated with smaller hippocampal volumes^7^, while other studies report no clear relationship^6,10^. Given this lack of consensus, we also investigated whether lesion size was independently associated with hippocampal volume, using a large sample of brain MRI scans with manually segmented stroke lesions.

The current study is a first step towards examining whether there is an association between the volume of the non-lesioned post-stroke hippocampus and sensorimotor impairment using a large cross-sectional dataset. In this study, we aimed to investigate the relationship between sensorimotor impairment and ipsilesional and contralesional hippocampal volumes (separately) in 357 participants with chronic stroke across 18 cohorts from the ENIGMA Stroke Recovery Working Group^29^. Due to the heterogeneity of post-stroke brain reorganization across individuals, large consortium-based multi-site studies such as the ENIGMA Stroke Recovery Working Group are important for achieving large and diverse samples that can identify associations that may have otherwise been undetectable in a smaller single-site sample^30^. In addition, the diversity of data allows us to verify whether associations are maintained beyond a single cohort, improving the robustness and generalizability of research findings. First, we investigated associations between sensorimotor impairment and hippocampal volume, controlling for lesion size and additional covariates of age, sex, and lesioned hemisphere. The Fugl-Meyer Assessment of Upper Extremity (FMA-UE) was used as a measure of sensorimotor impairment of the paretic upper limb^31^. We hypothesized that greater post-stroke sensorimotor impairment would be correlated with smaller ipsilesional but not contralesional hippocampal volume. Based on the involvement of the hippocampus in sensorimotor circuits, we hypothesized that the association between ipsilesional hippocampal volume and sensorimotor impairment would be independent of lesion size. Second, in an exploratory analysis, we tested to see if sex had a moderating effect on the relationship between sensorimotor impairment and hippocampal volume. Due to more severe hippocampal vulnerability and poorer stroke outcome trends in women, we hypothesized that women would have a stronger relationship between more severe sensorimotor impairment and smaller hippocampal volume than men. Finally, given the lack of consensus in the literature regarding lesion size and hippocampal volume, we independently tested associations between lesion size and hippocampal volume, without the FMA-UE included in the model. We hypothesized that larger lesion size would be significantly associated with smaller ipsilesional but not contralesional hippocampal volumes.

## Materials and methods

### 2.1 ENIGMA Stroke Recovery Dataset

A subset of cross-sectional data from the ENIGMA Stroke Recovery Working Group database (available as of December 15, 2020) was used. Details of the ENIGMA Stroke Recovery procedures and methods are available in Liew et al., 2020^29^. The data were collected across 18 research studies (cohorts) conducted at 10 different research institutes in six countries (**Table 1**).

**Table 1.** Demographics for ENIGMA Stroke Recovery Working Group participants included in the study by cohort. Total sample size (N), number of women and men, and information about age (years), Fugl-Meyer Assessment of Upper Extremity (FMA-UE), and raw lesion size in cubic centimeters (cc) are listed. For more information regarding cohort demographics by sex, see **Supplemental Table 1-2**.

| **Cohort** | **Total N (Women/Men)** | **Median Age (years)**<br>**(IQR, min-max)** | **Median FMA-UE**<br>**(IQR, min-max)** | **Median Lesion Size (cc)**<br>**(IQR, min-max)** |
| --- | --- | --- | --- | --- |
| Cohort 1 | 39 (10/29) | 61 (17, 31-80) | 43 (16, 0-58) | 6.1 (20.3, 0.04-120.8) |
| Cohort 2 | 12 (6/6) | 69.5 (12, 39-85) | 33 (27, 13-48) | 28.3 (28.5, 4.2-137.4) |
| Cohort 3 | 15 (6/9) | 61 (17, 33-85) | 16 (13, 5-40) | 21.1 (68.7, 0.6-182.2) |
| Cohort 4 | 19 (6/13) | 44 (15, 30-68) | 10 (11, 1-34) | 35.8 (54.4, 4.5-313.5) |
| Cohort 5 | 28 (12/16) | 64 (18, 44-81) | 52 (33, 8-65) | 1.9 (25.7, 0.1-237.7) |
| Cohort 6 | 10 (3/7) | 61 (12.5, 49-72) | 65 (3, 45-65) | 1.4 (1.1, 0.5-9.1) |
| Cohort 7 | 14 (5/9) | 58 (12, 45-69) | 63 (14, 6-65) | 2.0 (2.9, 0.04-6.9) |
| Cohort 8 | 11 (4/7) | 56 (12, 45-74) | 48 (15, 25-55) | 35.8 (50.2, 0.7-103.9) |
| Cohort 9 | 11 (3/8) | 59 (3, 45-68) | 38 (18, 15-49) | 2.6 (21.7, 0.7-53.7) |
| Cohort 10 | 8 (4/4) | 58 (8, 46-73) | 48 (16, 35-59) | 28.4 (43.2, 0.4-59) |
| Cohort 11 | 22 (6/16) | 61.5 (11, 23-75) | 49 (22, 23-64) | 5.6 (41.5, 0.4-201.4) |
| Cohort 12 | 13 (4/9) | 57 (13, 32-80) | 54 (15, 38-63) | 4.8 (18.2, 0.3-98) |
| Cohort 13 | 12 (4/8) | 66 (16, 31-83) | 51 (26, 19-62) | 4.4 (37.6, 0.2-107.5) |
| Cohort 14 | 29 (18/11) | 50 (15, 25-79) | 41 (13, 24-53) | 12.1 (28.6, 0.1-143.6) |
| Cohort 15 | 10 (3/7) | 61.5 (11, 42-76) | 29 (16, 11-60) | 9.1 (23.4, 3-186.1) |
| Cohort 16 | 40 (14/26) | 66.5 (11, 43-93) | 47 (30, 4-65) | 9.2 (26.1, 0.5-111.8) |
| Cohort 17 | 36 (15/21) | 70 (14, 37-80) | 53 (27, 8-65) | 7.6 (29.3, 0.3-188.4) |
| Cohort 18 | 28 (12/16) | 64 (14, 34-85) | 27 (5, 14-34) | 5 (29.4, 0.7-136.9) |
| **Total** | **357 (135/222)** | **61 (18, 23-93)** | **41 (28, 0-65)** | **7.6 (33.4, 0.04-313.5)** |

ENIGMA Stroke Recovery participants with the following data were included: 1) high resolution (1-mm isotropic) T1-weighted brain MRI (T1w) acquired with a 3T MRI scanner, 2) Fugl-Meyer Assessment of Upper Extremity score (FMA-UE; acquired on a scale from 0-66: 0 = severe sensorimotor impairment, 66 = no sensorimotor impairment), 3) age, and 4) sex. As we were interested in studying effects of secondary degeneration of the hippocampus, we only included participants with chronic stroke (defined as data acquired at least 180 days post-stroke^32^). Behavioral data were collected within approximately 72 hours of the MRI. Exclusion criteria included site-reported bilateral, brainstem, or cerebellar lesions, participants with no identifiable lesions, and participants with no sensorimotor impairment (FMA-UE = 66). In addition, each hippocampus was visually inspected with lesion masks overlaid, and any brains with hippocampal lesions were excluded. The total initial sample size was N = 357 (age: median = 61 years, interquartile range (IQR) = 18, range = 23-93; FMA-UE: median = 41, IQR = 28, range = 0-65; 135 women and 222 men) (**Table 1**).

### 2.2 MRI Data Analysis

Hippodeep, an automated convolutional neural network-based hippocampal segmentation algorithm, was used to segment ipsilesional and contralesional hippocampal volumes as well as estimated total head size from the T1 weighted MRI^33^. Hippodeep was previously found to be the most robust out of the freely available methods for segmenting the hippocampus in people with stroke pathology^34^. Hippocampal segmentations were visually inspected according to previously described protocols^29,34^. Any segmentations that were not properly segmented were marked as failed and excluded from the analysis. This resulted in different sample sizes for the ipsilesional and contralesional analyses. More information on demographics of samples after quality control can be found in **Supplemental Tables 1-2**. We performed a supplemental analysis using only participants with hippocampal segmentations that passed quality control for both ipsilesional and contralesional hippocampi and confirmed that differences in sample sizes did not significantly influence the results (see **Supplemental Tables 3-6**). To account for differences in head size, hippocampal volume was normalized for head size by taking the ratio of hippocampal volume to head size for each participant and multiplying it by the average head size across the sample, as done in previous studies of post-stroke hippocampal volume^7,10,35^.

### 2.3 Manually Segmented Lesions

Lesions were manually segmented on the T1w MRI by B.L., M.D., J.S., A.Z.P., and S-L.L. according to an updated version of the Anatomical Tracings of Lesions After Stroke (ATLAS) protocol^36^. Briefly, brain lesions were identified, and masks were manually drawn on each individual brain in native space using ITK-Snap^37^. Each lesion was checked for quality by at least two different tracers. An expert neuroradiologist (G.B.) was also consulted to ensure lesion segmentation accuracy.

Although all participants were listed by the providing research sites as having unilateral lesions, additional secondary lesions were discovered in 100 participants during manual tracing, which were likely silent, subclinical, and/or prior strokes. Secondary lesions were found in both hemispheres, the brainstem, and the cerebellum, and ranged in size. For this paper, we refer to the primary lesioned hemisphere as the lesioned hemisphere noted by the research site. We also performed follow-up analyses excluding participants with any identified secondary lesions, which did not significantly impact results (**Supplementary Materials**). Lesion probability maps were generated by nonlinearly normalizing lesion masks and registering them to the MNI-152 template **(Figure 1)**.

**Figure 1.**
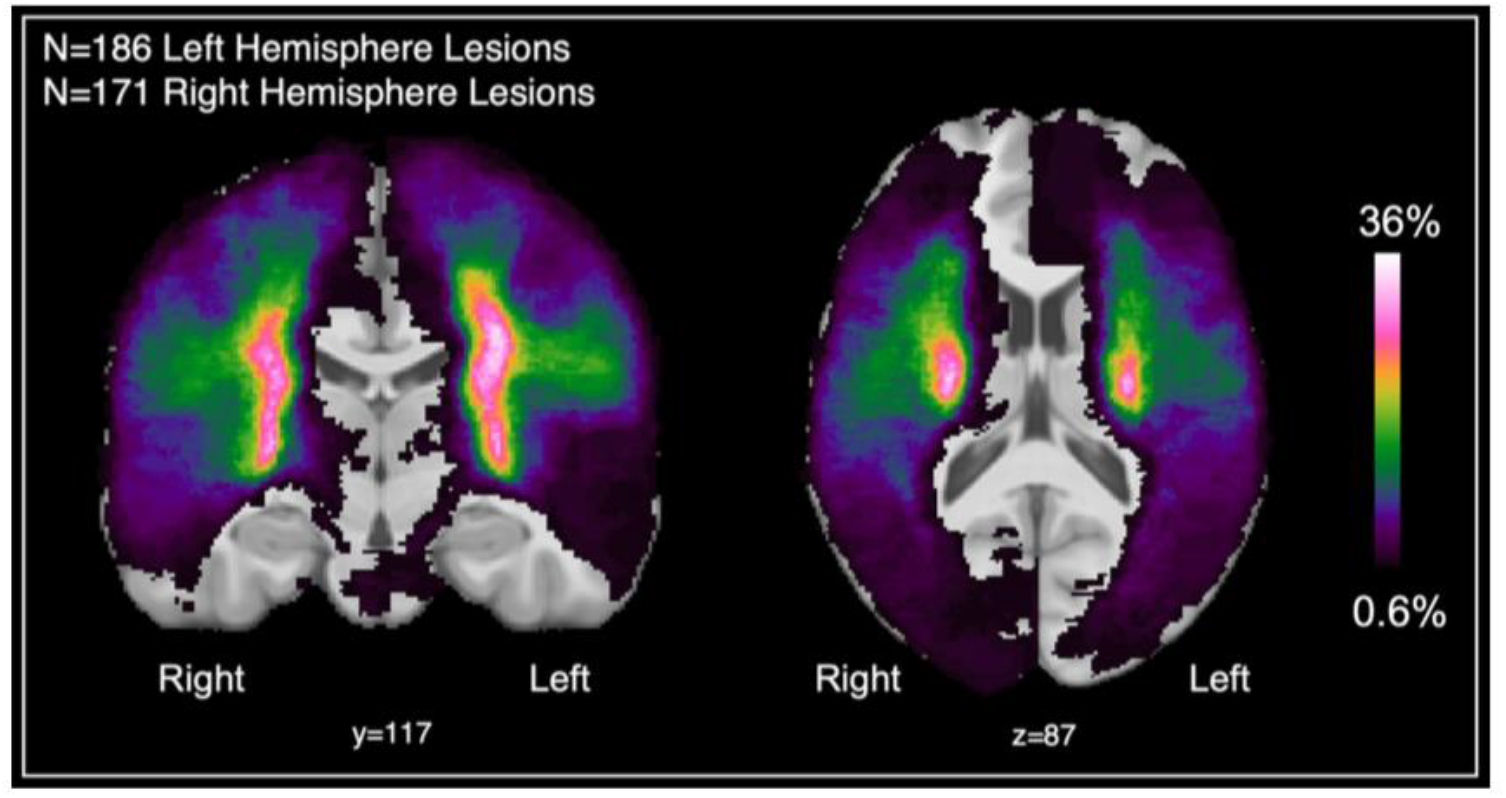
Lesion density maps for primary lesions from participants with cohort-reported left and right hemisphere lesions are overlaid on the MNI-152 template. Lesioned hemisphere refers to the primary lesion, as reported by the research cohort. The color bar refers to the percentage of overlapping lesions across participants.

Finally, lesion volume was calculated by summing the voxels within each manually traced lesion mask. Lesion size was also normalized for head size as previously described for hippocampal volume in *Methods Section 2.2*. Lesion size was then log transformed to normalize the distribution of the data.

### 2.4 Statistical Analysis

#### 2.4.1 Hippocampal Volume and Sensorimotor Impairment

We first tested our primary hypothesis that more severe post-stroke sensorimotor impairment is correlated with smaller ipsilesional but not contralesional hippocampal volumes by performing robust mixed-effects linear regressions with hippocampal volume as the dependent variable (see *Model 1*). Sensorimotor impairment was measured using the FMA-UE^31^. Sex (coded as a binary variable: women = 0, men = 1), age, and lesioned hemisphere (coded as binary variable: left hemisphere lesion = 0.5, right hemisphere lesion = 1.5) were included in the model as fixed effects and cohort was included in the model as a random effect:

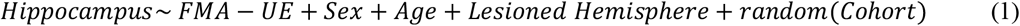

Next, we tested our exploratory hypothesis that sex may have a moderating effect on the relationship between sensorimotor impairment and hippocampal volume by including an FMA-UE*Sex interaction covariate as a fixed effect (*Model 2):*

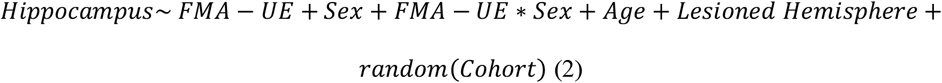

Sex differences in sensorimotor impairment, lesion size, and age were tested using an independent *t*-test.

Finally, we tested our hypothesis that sensorimotor impairment is independently associated with hippocampal volume regardless of lesion size by including lesion size as a fixed covariate *(Model 3)*:

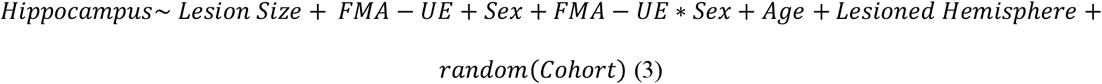

#### 2.4.2 Associations between Lesion Size and Hippocampal Volume

To investigate associations between lesion size and hippocampal volume, regardless of sensorimotor impairment, we performed a robust mixed effects regression with ipsilesional and contralesional hippocampal volume as dependent variables with lesion size, age, sex, and lesioned hemisphere as fixed effects, and cohort as a random effect (*Model 4)*:

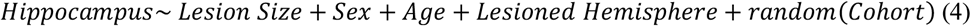

### 2.5 Statistical Tools

All statistical analyses were performed in R (version 4.0.2^38^). The Mahalanobis distance was used to detect multivariate outliers^39^, which were then removed from the analyses (**Supplemental Materials**). All continuous measures were normalized using the *scale* function in R to be analyzed as z-scores to calculate standardized beta coefficients. All mixed effects regressions were initially run as linear mixed effects regressions (*lmer* function from *nlme* package). Collinearity for variables in every model tested was ruled out (variance inflation factor ≤ 2.5). Regression assumptions of linearity, normality of the residuals, and homogeneity of the residual variance were tested by visually inspecting residuals versus fits plots as well as qq-plots. After detecting influential observations using Cook’s distance in each analysis^40^, the analyses were repeated using robust mixed-effects regression. Robust mixed effects regression (*rlmer* from the *robustlmm* package) avoids excluding data by reducing the weight of influential observations^41^. We therefore report the results of the robust mixed effects regression. For all analyses, beta coefficients for the factor of interest and confidence intervals (Beta(CI)), standard error (SE), t-value and degrees of freedom (t(DF)), standardized effect size (d-value), and uncorrected *p*-values are reported. For each analysis, a Bonferroni correction was applied for two comparisons (ipsilesional, contralesional; corrected *p*-value < 0.025). Effect sizes were mapped onto a template of the hippocampus to visualize the results using *ggseg*^42^.

#### Data availability

To protect the privacy of research participants, individual subject data used in this study is not available in a public repository. Participating research cohorts vary in public data sharing restrictions as determined by a) ethical review board and consent documents; b) national and transnational sharing laws; and c) institutional processes that may require signed data transfer agreements for limited, predefined data use. However, data sharing is possible for new and participating ENIGMA Stroke Recovery working group members who agree to the consortium’s ethical standards for data use and upon the submission of a secondary analysis plan for group review. Upon the approval of the proposed analysis plan, access to relevant data is provided contingent on local PI approval, data availability, and compliance with supervening regulatory boards. Deidentified summary data as well as code used for this study can be made available upon reasonable request by the corresponding author.

## Results

### 3.1 Hippocampal Volume and Sensorimotor Impairment

Greater sensorimotor impairment was significantly associated with smaller ipsilesional (*Beta* = 0.16, *p*-*value* = 0.005, *R*^*2*^ = 0.27) but not contralesional (*Beta* = 0.003, *p*-*value* = 0.96, *R*^*2*^ = 0.29) hippocampal volume after adjusting for age, sex, lesioned hemisphere, and cohort (**Table 2**). When including FMA-UE*Sex interaction as a covariate, we observed a better model fit and an increase in effect size for the association between sensorimotor impairment and ipsilesional hippocampal volume (*Beta* = 0.31, *p*-*value* < 0.001, *R*^*2*^ = 0.30; **Table 3)**. Furthermore, FMA-UE remained independently associated with ipsilesional hippocampal volume after including lesion size in the model (*Beta* = 0.26*, p*-*value* = 0.001, *R*^*2*^ = 0.33; **Table 4, Figure 2**). This association remained significant when excluding participants with secondary lesions (*Beta* = 0.30, *p*-*value* = 0.001, *R*^*2*^ = 0.35; **Supplemental Table 7.**).

**Table 2.** Summary statistics from robust mixed-effects linear regression to test associations between ipsilesional hippocampal volume and sensorimotor impairment (*top*) and contralesional hippocampal volume and sensorimotor impairment (*bottom*). The full model as well as the sample size (*N*), conditional R^2^, beta coefficient (*Beta*) with 95% confidence interval (*CI*), standard error (*SE*), *t*-*value* and degrees of freedom *t*(*DF*), standardized *d*-*value*, uncorrected *p*-*value* for all fixed effect covariates are reported. Significant covariates are denoted in bold.

| *Hippocampus ~ FMA-UE + Sex + Lesioned Hemisphere + Age + random(Cohort)* |  |  |  |  |  |
| --- | --- | --- | --- | --- | --- |
| *Covariates* | *Beta(CI)* | *SE* | *t(DF)* | *d-value* | *p-value* |
| ***IPSILESIONAL HIPPOCAMPAL VOLUME (N=336; R<sup>2</sup>=0.27)*** |  |  |  |  |  |
| **FMA-UE** | 0.16 (0.05 – 0.27) | 0.06 | 2.80(287) | 0.33 | **0.005** |
| **Sex** | -0.53 (-0.73 – -0.33) | 0.10 | -5.21(324) | -0.58 | **<0.001** |
| Lesioned Hemisphere | 0.19 (-0.01 – 0.39) | 0.10 | 1.84(336) | 0.20 | 0.06 |
| **Age** | -0.32 (-0.42 – -0.22) | 0.05 | -6.16(335) | -0.67 | **<0.001** |
| ***CONTRALESIONAL HIPPOCAMPAL VOLUME (N=349; R<sup>2</sup>=0.29)*** |  |  |  |  |  |
| FMA-UE | 0.003 (-0.10 – 0.11) | 0.05 | 0.05(238) | 0.01 | 0.96 |
| **Sex** | -0.50 (-0.69 – -0.31) | 0.10 | -5.14(343) | -0.56 | **<0.001** |
| **Lesioned Hemisphere** | -0.32 (-0.51 – -0.13) | 0.10 | -3.30(346) | -0.35 | **0.001** |
| **Age** | -0.41 (-0.51 – -0.32) | 0.05 | -8.30(346) | -0.89 | **<0.001** |

**Table 3.** Summary statistics from robust mixed-effects linear regression to test associations between ipsilesional hippocampal volume and sensorimotor impairment (*top*) and contralesional hippocampal volume and sensorimotor impairment (*bottom*) when including a sensorimotor impairment and sex interaction. The full model as well as the sample size (*N*), conditional R^2^, beta coefficient (*Beta*) with 95% confidence interval (*CI*), standard error (*SE*), *t*-*value* and degrees of freedom *t*(*DF*), standardized *d*-*value*, uncorrected *p*-*value* for all fixed effect covariates are reported. Significant covariates are denoted in bold.

| *Hippocampus ~ FMA-UE\*Sex + FMA-UE + Sex + Lesioned Hemisphere + Age + random(Cohort)* |  |  |  |  |  |
| --- | --- | --- | --- | --- | --- |
| *Covariates* | *Beta(CI)* | *SE* | *t*(*DF*) | *d*-value | *p*-value |
| ***IPSILESIONAL HIPPOCAMPAL VOLUME (N=336; R<sup>2</sup>=0.30)*** |  |  |  |  |  |
| **FMA-UE** | 0.31 (0.15 – 0.46) | 0.08 | 3.86(336) | 0.42 | **<0.001** |
| **FMA-UE\*Sex** | -0.26 (-0.46 – -0.07) | 0.10 | -2.61(324) | -0.29 | **0.009** |
| **Sex** | -0.53 (-0.73 – -0.33) | 0.10 | -5.29(336) | -0.58 | **<0.001** |
| Lesioned Hemisphere | 0.17 (-0.03 – 0.37) | 0.10 | 1.69(335) | 0.18 | 0.09 |
| **Age** | -0.32 (-0.42 – -0.22) | 0.05 | -6.27(332) | -0.69 | **<0.001** |
| ***CONTRALESIONAL HIPPOCAMPAL VOLUME (N=349; R<sup>2</sup>=0.32)*** |  |  |  |  |  |
| FMA-UE | 0.16 (0.01 – 0.31) | 0.08 | 2.06(334) | 0.23 | 0.04 |
| **FMA-UE\*Sex** | -0.27 (-0.46 – -0.08) | 0.10 | -2.76(348) | -0.30 | **0.006** |
| **Sex** | -0.51 (-0.70 – -0.32) | 0.10 | -5.28(343) | -0.57 | **<0.001** |
| **Lesioned Hemisphere** | -0.35 (-0.54 – -0.16) | 0.10 | -3.58(343) | -0.39 | **<0.001** |
| **Age** | -0.41 (-0.51 – -0.32) | 0.05 | -8.38(344) | -0.90 | **<0.001** |

**Table 4.**
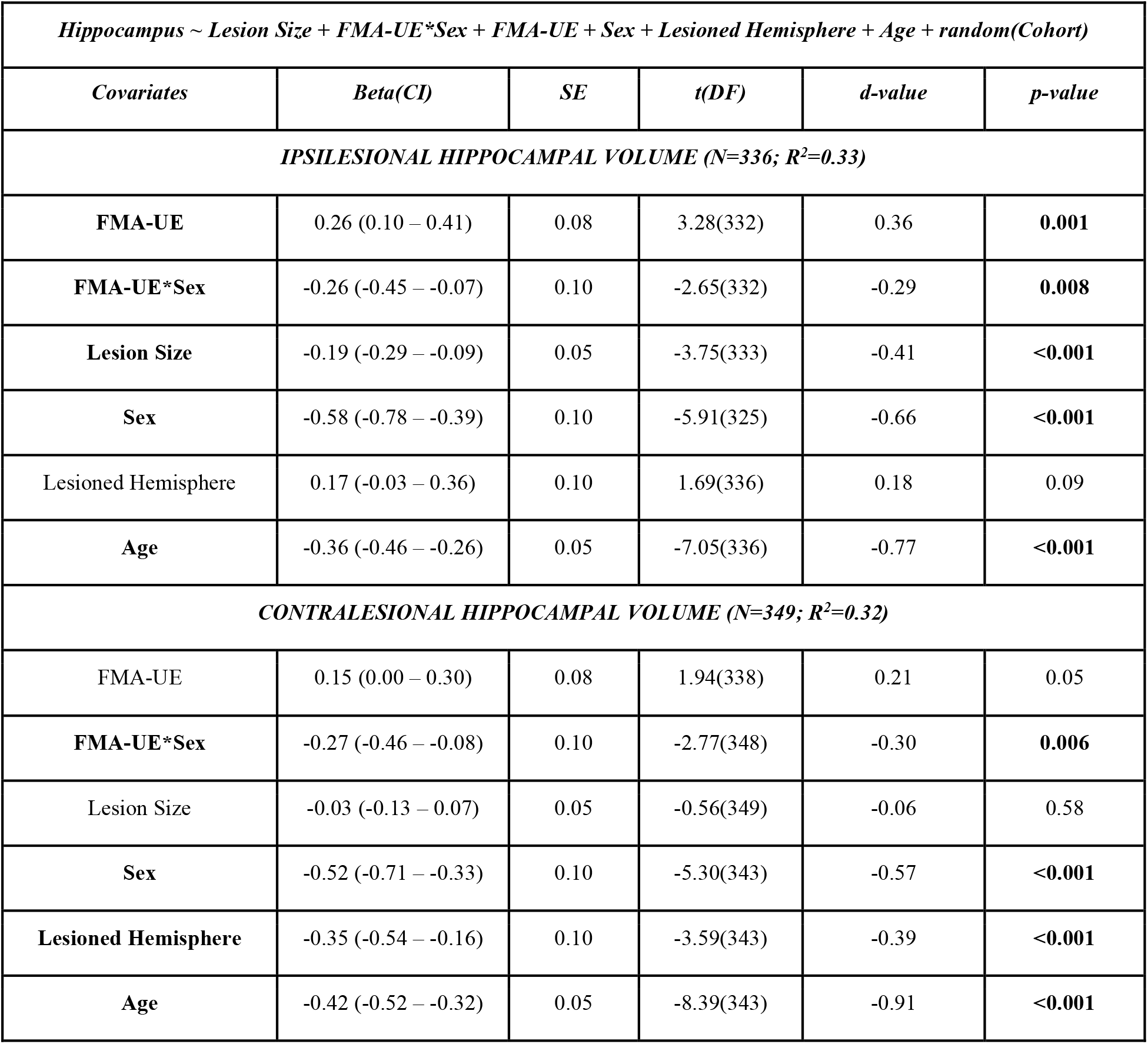
Summary statistics from robust mixed-effects linear regression to test associations between ipsilesional hippocampal volume and sensorimotor impairment (*top*) and contralesional hippocampal volume and sensorimotor impairment (*bottom*) when including lesion size as a covariate. The full model as well as the sample size (*N*), conditional R^2^, beta coefficient (*Beta*) with 95% confidence interval (*CI*), standard error (*SE*), *t*-*value* and degrees of freedom *t*(*DF*), standardized *d*-*value*, uncorrected *p*-*value* for all fixed effect covariates are reported. Significant covariates are denoted in bold.

**Figure 2.**
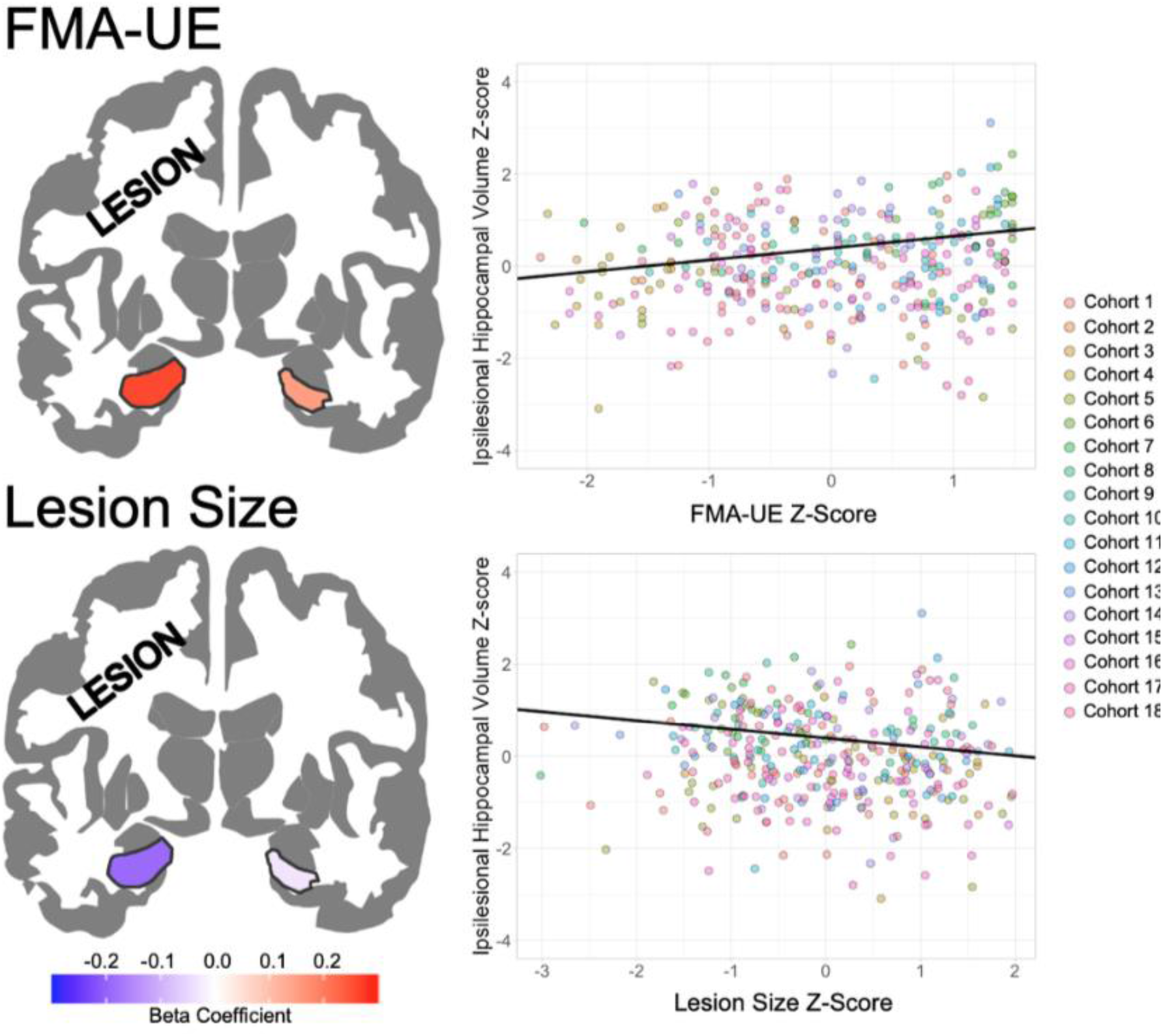
Effect sizes (standardized *Beta* values) for ipsilesional and contralesional hippocampi are mapped onto a template for associations between hippocampal volumes and sensorimotor impairment (*top left*) and lesion size (*bottom left*), with warmer colors representing stronger positive associations. Trend lines (*black line*) are plotted for the association between ipsilesional hippocampal volume z-scores with FMA-UE z-scores (*top right*) and lesion size z-scores (*bottom right*). Scatter plot points are colored by research cohort.

### 3.2 Sex Effects on the Association between Hippocampal Volume and Sensorimotor Impairment

A *t*-test revealed no significant differences in FMA-UE (*t*(260) = 1.13, *p*-*value* = 0.26) or age (*t*(249) = 1.12, *p*-*value* = 0.26) between women and men. Women did have significantly larger lesions than men (*t*(277) = 2.9, *p*-*value* = 0.004) (**Figure 3**). The FMA-UE*Sex interaction was a significant covariate for both ipsilesional (*Beta* = −0.26, *p*-*value* = 0.009, *R*^*2*^ = 0.30) and contralesional (*Beta* = −0.27, *p*-*value* = 0.006, *R*^*2*^ = 0.32) hippocampal volumes (**Figure 3, Table 3**), even after accounting for lesion size (ipsilesional: *Beta* = −0.26, *p*-*value* = 0.008, *R*^*2*^ = 0.33; contralesional: *Beta* = −0.27, *p*-*value* = 0.006, *R*^*2*^ = 0.32; **Table 4**). In the ipsilesional hippocampus, women had a positive slope (*β* = 0.26) and men had a negative slope close to 0 (*β* = −0.002). In the contralesional hippocampus, women had a positive slope (*β* = 0.15) while men had a negative slope (*β* = −0.12) **(Figure 3**). The FMA-UE*Sex interaction remained significantly associated with both ipsilesional (*Beta* = −0.31, *p*-*value* = 0.008, *R*^*2*^ = 0.35) and contralesional (*Beta* = −0.28, *p*-*value* = 0.017, *R*^*2*^ = 0.32) hippocampal volumes, even when excluding participants with secondary lesions **(Supplemental Table 7)**.

**Figure 3.**
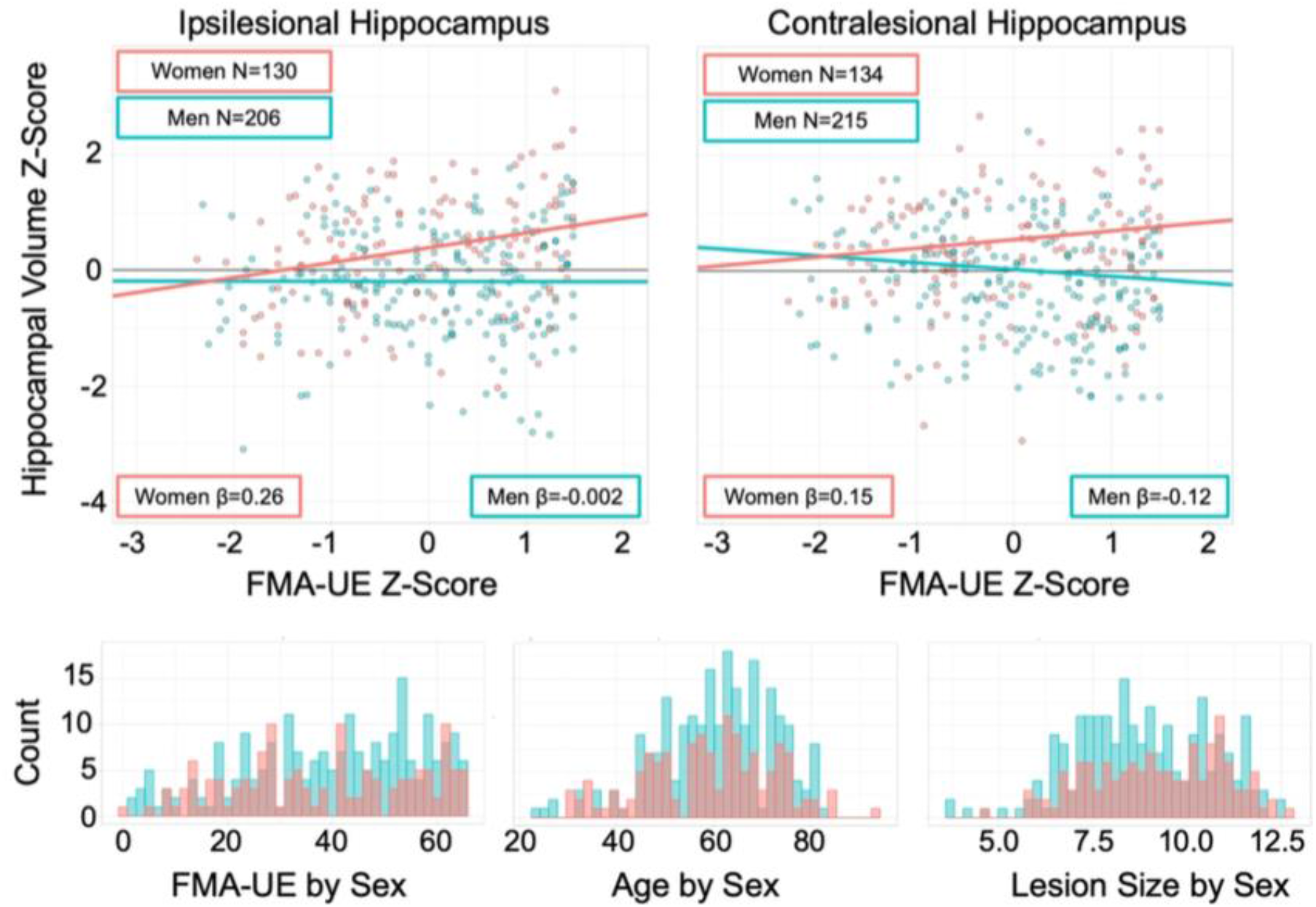
Trend lines are plotted for the association between FMA-UE z-score (x-axis) and hippocampal volumes z-score (y-axis) for women (*red*) and men (*blue*) calculated from the FMA-UE*Sex interactions. Histograms for FMA-UE scores (*bottom left*), age (*bottom middle*), and lesion size (*bottom right*) are plotted by sex (women in *red* and men in *blue*).

### 3.3 Hippocampal Volume and Lesion Size

Larger lesion size was significantly associated with smaller ipsilesional (*Beta* = −0.21, *p*-*value* < 0.001, *R*^*2*^ = 0.33) but not contralesional hippocampal volume (*Beta* = −0.03, *p*-*value* = 0.60, *R*^*2*^ = 0.30), after adjusting for age, sex, lesioned hemisphere, and cohort (**Table 5; Figure 2**). Lesion size remained significantly associated with smaller ipsilesional, but not contralesional, hippocampal volume, even when excluding participants with secondary lesions (*Beta* = −0.18, *p*-*value* = 0.003, *R*^*2*^ = 0.33; **Supplemental Table 8**).

**Table 5.** Summary statistics from robust mixed-effects linear regression to test associations between ipsilesional hippocampal volume and lesion size (*top*) and contralesional hippocampal volume and lesion size (*bottom*). The full model as well as the sample size (*N*), conditional R^2^, beta coefficient (*Beta*) with 95% confidence interval (*CI*), standard error (*SE*), *t*-*value* and degrees of freedom *t*(*DF*), standardized *d*-*value*, uncorrected *p*-*value* for all fixed effect covariates are reported. Significant covariates are denoted in bold.

| *Hippocampus ~ Lesion Size + Sex + Lesioned Hemisphere + Age + random(Cohort)* |  |  |  |  |  |
| --- | --- | --- | --- | --- | --- |
| *Covariates* | *Beta(CI)* | *SE* | *t(DF)* | *d-value* | *p-value* |
| ***IPSILESIONAL HIPPOCAMPAL VOLUME (N=336; R<sup>2</sup>=0.33)*** |  |  |  |  |  |
| **Lesion Size** | -0.21 (-0.31 – -0.12) | 0.05 | -4.23(334) | -0.46 | **<0.001** |
| **Sex** | -0.58 (-0.77 – -0.38) | 0.10 | -5.78(324) | -0.64 | **<0.001** |
| Lesioned Hemisphere | 0.16 (-0.03 – 0.36) | 0.10 | 1.64(336) | 0.18 | 0.10 |
| **Age** | -0.35 (-0.45 – -0.25) | 0.05 | -6.83(335) | -0.75 | **<0.001** |
| ***CONTRALESIONAL HIPPOCAMPAL VOLUME (N=349; R<sup>2</sup>=0.30)*** |  |  |  |  |  |
| Lesion Size | -0.03 (-0.12 – 0.07) | 0.05 | -0.53(348) | -0.06 | 0.60 |
| **Sex** | -0.51 (-0.70 – -0.32) | 0.10 | -5.17(343) | -0.56 | **<0.001** |
| **Lesioned Hemisphere** | -0.32 (-0.51 – -0.13) | 0.10 | -3.34(346) | -0.36 | **0.001** |
| **Age** | -0.42 (-0.52 – -0.32) | 0.05 | -8.36(344) | -0.90 | **<0.001** |

## Discussion

In this study, associations between sensorimotor impairment, lesion size, sex and hippocampal volume were investigated in participants with chronic stroke from 18 research cohorts in the ENIGMA Stroke Recovery working group^29^. Greater sensorimotor impairment and larger lesion sizes were both significantly associated with smaller ipsilesional hippocampal volumes, and the association between sensorimotor impairment and hippocampal volume was stronger in women than in men.

To our knowledge, this is the first study to report associations between hippocampal volume and sensorimotor impairment in chronic stroke patients. Greater sensorimotor impairment was independently associated with smaller ipsilesional hippocampal volume, even after adjusting for lesion size. This suggests that post-stroke ipsilesional hippocampal integrity may be related to sensorimotor impairment. Spreading depression (SD) might explain secondary degeneration of the ipsilesional hippocampus, where neurotoxic signals from the core of the lesion propagate to adjacent grey matter regions and cause damage^6,7,10^. The ionic imbalance that results from a blockage of blood supply during the acute phase of a stroke causes a buildup of extracellular glutamate that leads to a self-propagating wave of cell depolarization throughout neighboring gray matter^43^. The hippocampus is filled with tightly packed, easily excitable glutamatergic neurons and a high density of N-methyl-D-aspartate receptors^44^, making it more susceptible to damage from SD. Overexcitation of the hippocampal glutamatergic network leads to hippocampal excitotoxicity, resulting in hippocampal neuron apoptosis, which is thought to be reflected on a macroscale as reduced hippocampal volume^10^. The damaging effects of SD are likely more prominent in the lesioned hemisphere because SD waves do not propagate easily through white matter^45^, therefore the waves cannot easily traverse to the contralesional hippocampus. While a magnetic resonance spectroscopy study reported evidence of hippocampal neuronal loss in the contralesional hippocampus, it was less severe and not detectable using volumetric MRI^10^. SD is still not well understood, therefore contralesional hippocampal damage may be caused by mild SD or might be attributed to other forms of secondary degeneration such as diaschisis^5^. The available evidence is insufficient to support SD as the key cause of reduced ipsilesional hippocampal volumes observed in this study, however these findings could provide future directions for research investigating the mechanisms of stroke-related hippocampal damage.

In addition to hippocampal damage incurred by SD, ipsilesional disruption to sensorimotor circuits may cause secondary degeneration of the ipsilesional hippocampus, possibly as a result of anatomical connectivity to damaged areas (e.g., through the thalamus^20^, basal ganglia^15^, sensorimotor cortex^18^, or supplementary motor area^21^) via anterograde degeneration. Furthermore, as the hippocampus is an important limbic system structure, it is heavily involved in learning, memory, and emotion^46^. Post-stroke cognitive impairment^47,48^, depression^47^, and anxiety^49^ are all common pervasive symptoms in stroke survivors that interfere with rehabilitation and are associated with poor stroke outcomes^47,48^. Limbic system disruption caused by secondary post-stroke hippocampal damage may cause cognitive impairment or aggravate symptoms of depression and anxiety, which in turn, may interfere with stroke sensorimotor rehabilitation efforts. Further functional and longitudinal research is necessary to understand the relationship between hippocampal damage and sensorimotor circuits and how hippocampal volume loss may impact sensorimotor rehabilitation.

In an exploratory analysis, we found significant sex differences in the association between FMA-UE and bilateral hippocampal volume, where women showed progressively greater sensorimotor impairment with smaller hippocampal volumes compared to men. This observation suggests that women with greater sensorimotor impairment may also have more hippocampal damage or more pre-existing hippocampal atrophy compared to men. In addition, sex differences observed in the association between sensorimotor impairment and hippocampal volume did not appear to be driven by age or severity of sensorimotor impairment. Although lesion size was significantly larger in women, the FMA-UE*Sex interaction covariate was independently associated with hippocampal volume, even when accounting for lesion size. Overall, these findings should be considered exploratory given the unequal number of men and women in the sample. Further research is needed to confirm these findings, as our sample was unable to account for additional variables^50,51^ thought to influence the hippocampus in a sex-dependent way such as estrogen levels^24^, dementia^52–54^, and depression^55^. Furthermore, the extent to which sex differences observed in stroke research are a result of physiological differences between sexes versus different contextual factors such as treatment received by women post-stroke remains unclear^27,56^. Further research on sex differences in stroke is crucial to improve our understanding of the relationship between hippocampal damage and sensorimotor impairment.

Lastly, we found that larger lesion sizes were significantly associated with smaller hippocampal volumes, but only within the lesioned hemisphere, independent of sensorimotor impairment. This finding is in line with a previous study^7^ and may indicate that smaller hippocampal volumes observed in stroke patients may be specific to the amount of stroke-related damage within the lesioned hemisphere beyond that which is attributed to age-related atrophy^46^ or other stroke risk factors such as hypertension^57^ or changes in estrogen^24^ that are typically observed bilaterally.

### Limitations and Future Directions

This study only considered gross hippocampal volume. However, the hippocampus is composed of structurally and functionally distinct subfields, each differentially vulnerable to disease^46,58^. Structurally, reduced neuron density has been observed in the cells of the CA1 but not CA2 subfield of post-mortem stroke patients when compared to controls^59^, and larger white matter hyperintensity volume has been associated with reduced volume of the hippocampal-amygdala transition area^60^. Functionally, while the posterior extents of the hippocampus along the long axis are thought to be more involved with memory and cognitive processing^61^, the anterior extents have been implicated in sensorimotor integration^46^. Further research investigating sensorimotor impairment and the hippocampus at a finer resolution, such as at the level of hippocampal subfields^58^ or vertex-wise associations^62^, may reveal more specific and robust relationships that can better inform the understanding the impact of hippocampal damage on recovery and rehabilitation.

In addition, although secondary lesions were discovered while manually tracing lesion masks, our findings did not change when participants with secondary lesions were excluded. Further research is necessary to investigate the impact of lesion location on the association between hippocampal volume and sensorimotor impairment.

Given the focus on hippocampal volumes, another limitation of this study is the lack of cognitive and depression data. While cognitive and depressive scores are available for a small number of cohorts in the ENIGMA Stroke Recovery database, the participants with available data have very limited information. Many of the participating stroke sensorimotor rehabilitation research studies also used cognitive impairment as an exclusion criteria^63^, resulting in participants with no or mild cognitive deficits.

Finally, the current sample is cross-sectional and cannot account for the extent of longitudinal hippocampal atrophy that may have occurred as a result of stroke, mild cognitive impairment, pre-existing dementia, or normal aging. This sample also does not contain data on type of dose of rehabilitation treatment received, which could also influence sensorimotor outcomes. However, the current cross-sectional analysis serves as a first step to examining the relationship between hippocampal volumes, sensorimotor impairment, lesion volume and sex and can be used to guide future questions using a longitudinal dataset.

### Conclusion

Our findings demonstrate a novel association between chronic post-stroke sensorimotor impairment and hippocampal volume that may be modulated by sex. We provide supporting evidence to existing literature that reduced hippocampal volume is likely a consequence of stroke-related damage within the lesioned hemisphere. Overall, these findings provide unique insight into the role that the hippocampus may play in post-stroke sensorimotor impairment.

## Acknowledgements

We thank all of the members of the ENIGMA Stroke Recovery Working Group and all of the participants and their families for their contribution to this study. We also thank Sophia Thomopoulos for her assistance.

## Abbreviations

ENIGMA: Enhancing NeuroImaging Genetics through Meta-Analysis
FMA-UE: Fugle-Meyer Assessment of Upper Extremity
IQR: Interquartile range
SD: Spreading depression

