## Supplemental Material for "Chronic stroke sensorimotor impairment is related to smaller hippocampal volumes: An ENIGMA analysis"

### Supplementary Materials

#### *Data demographics following removal of multivariate outliers and segmentations that failed quality control*

3 participants were identified as multivariate outliers using the Mahalanobis distance. These participants were excluded from the analysis. Additionally, 18 participants were excluded from the ipsilesional analysis (**Supplemental Table 1**) and 5 participants were excluded from the contralesional analysis (**Supplemental Table 2**) due to hippocampal segmentations that failed quality control. More details regarding ENIGMA quality control protocols can be found in Zavaliangos-Petropulu et al., 2020 and Liew et al., 2020. Briefly, a segmentation failed quality control if the segmentation grossly underestimated the hippocampus, overestimated the hippocampus by including regions of the brain outside of the hippocampus, or missed the hippocampus entirely. To ensure that differing sample sizes for the ipsilesional and contralesional analyses did not influence the results, we performed a supplemental analysis only including participants with hippocampal segmentations that passed quality control for both ipsilesional and contralesional hippocampi (**Supplemental Tables 3-6)**.

### *Supplementary Tables*

**Supplemental Table 1.** Detailed demographics for the sample used to analyze sensorimotor impairment and ipsilesional hippocampal volume. Demographics for women and men are broken down by cohort. Sample size (N) and median (IQR, range) of age, FMA-UE, and raw lesion size are reported.

| | **Women** | | | | **Men** | | | |
|---|---|---|---|---|---|---|---|---|
| **Cohort** | **N** | **Age (yrs)** | **FMA-UE** | **Lesion Size (cc)** | **N** | **Age (yrs)** | **FMA-UE** | **Lesion Size (cc)** |
| Cohort 1 | 10 | 53 (14,31-75) | 35 (16,0-58) | 8.1 (31.5,1.1-96.6) | 26 | 65 (12,42-80) | 44 (13,19-56) | 3.8 (7.6,0-37) |
| Cohort 2 | 3 | 64 (17,51-85) | 31 (17,14-48) | 9.8 (24.6,6.3-55.5) | 5 | 71 (18,39-74) | 37 (22,13-46) | 10 (23.4,4.2-34.1) |
| Cohort 3 | 6 | 62 (9,35-85) | 15 (7,5-18) | 21.6 (16.1,1.3-112.4) | 6 | 52 (15,33-71) | 29 (11,5-40) | 7.6 (6.7,0.6-113.4) |
| Cohort 4 | 5 | 34 (19,30-68) | 11 (5,9-34) | 48.8 (23.4,9.7-90) | 10 | 46 (11,32-63) | 9 (11,1-24) | 31.9 (30,8.1-130.2) |
| Cohort 5 | 12 | 62.5 (14,44-81) | 55 (21,22-65) | 1.7 (6.2,0.1-70.2) | 16 | 67 (18,50-81) | 40 (32,8-65) | 5.2 (66.6,0.3-237.7) |
| Cohort 6 | 3 | 63 (10, 2-72) | 65 (0,65-65) | 5 (4.3,0.5-9.1) | 7 | 59 (14,49-66) | 64 (9,45-65) | 1.3 (0.5,0.6-2.2) |
| Cohort 7 | 5 | 61 (4,57-65) | 49 (43,14-63) | 4.3 (1.3,0.3-6.9) | 9 | 53 (10,45-69) | 63 (4,6-65) | 1.5 (1.2,0.04-5.3) |
| Cohort 8 | 4 | 58 (16,45-74) | 50 (10,32-55) | 45.2 (43,0.7-64.6) | 7 | 56 (9,45-68) | 45 (15,25-55) | 35.8 (47.2,0.9-103.9) |
| Cohort 9 | 3 | 59 (1, 8-60) | 42 (11,26-49) | 2.5 (11.6,0.8-24) | 8 | 59 (5,45-68) | 36 (20,15-49) | 4.9 (22,0.7-53.7) |
| Cohort 10 | 4 | 61 (11,46-73) | 55 (9,35-58) | 10.9 (28,0.4-59) | 3 | 58 (4,56-64) | 43 (11,37-59) | 37 (27.3,1.4-55.9) |
| Cohort 11 | 6 | 65 (5,51-75) | 47 (24,23-61) | 5.2 (31.3,0.4-201.4) | 16 | 59 (10,23-75) | 49 (18,29-64) | 5.8 (42,0.6-71) |
| Cohort 12 | 4 | 51 (16,32-62) | 58 (8,54-63) | 1.8 (14.5,0.3-55.9) | 9 | 58 (7,47-80) | 48 (11,38-62) | 8.2 (17.8,1.3-98) |
| Cohort 13 | 4 | 68 (20,31-75) | 60 (4,51-62) | 23.2 (55,0.5-107.5) | 7 | 68 (19,52-83) | 37 (31,19-61) | 4.5 (30.3,0.2-62.6) |
| Cohort 14 | 17 | 50 (15,36-79) | 41 (15,24-47) | 13.6 (44.7,0.7-143.6) | 11 | 50 (15,25-76) | 44 (9,31-53) | 3.8 (12.5,0.1-32.4) |
| Cohort 15 | 3 | 47 (11,42-63) | 18 (5,11-21) | 36.7 (91.4,3.2-186.1) | 7 | 62 (12,51-76) | 35 (16,23-60) | 8.6 (3.3,3-97.5) |
| Cohort 16 | 14 | 68 (12,43-93) | 48 (33,20-65) | 12.1 (24.9,1.1-54.4) | 26 | 65 (9,45-81) | 47 (26,4-62) | 8 (25.2,0.5-111.8) |
| Cohort 17 | 15 | 68 (19,37-79) | 38 (34,8-64) | 7.6 (31.2,0.5-188.4) | 20 | 72 (12,51-80) | 56 (11,23-65) | 5.2 (22,0.3-110.8) |
| Cohort 18 | 12 | 62 (15,34-85) | 28 (5,14-34) | 15.6 (47.9,0.8-136.9) | 13 | 65 (17,50-78) | 27 (4,23-33) | 3.1 (3.6,0.7-34.1) |
| **Total** | **130** | **60 (20,30-93)** | **39 (29,0-65)** | **9.3 (39.5,0.11-201.4)** | **206** | **62 (16,23-83)** | **44 (25,1-65)** | **5.6 (24.1,0.04-237.7)** |

**Supplemental Table 2.** Detailed demographics for the sample used to analyze sensorimotor impairment and contralesional hippocampal volume. Demographics for women, men, and total are broken down by cohort. Sample size (N) and median (IQR, range) of age, FMA-UE, and raw lesion size are reported.

| | Women | | | | Men | | | |
|---|---|---|---|---|---|---|---|---|
| **Cohort** | **N** | **Age (yrs)** | **FMA-UE** | **Lesion Size (cc)** | **N** | **Age (yrs)** | **FMA-UE** | **Lesion Size (cc)** |
| Cohort 1 | 10 | 53 (14,31-75) | 35 (16,0-58) | 8.1 (31.5,1.1-96.6) | 28 | 64 (17,32-80) | 44 (13,19-56) | 4.9 (12,0.04-120.8) |
| Cohort 2 | 6 | 66 (9,51-85) | 23 (29,13-48) | 38 (39,6.3-137.4) | 6 | 71 (14,39-74) | 36 (17,13-46) | 19 (26.2,4.2-34.1) |
| Cohort 3 | 6 | 62 (9,35-85) | 15 (7,5-18) | 21.6 (16.1,1.3-112.4) | 8 | 57 (17,33-71) | 22 (22,5-40) | 10.5 (50.6,0.6-171.2) |
| Cohort 4 | 5 | 34 (19,30-68) | 11 (5,9-34) | 48.8 (23.4,9.7-90) | 12 | 45 (11,32-63) | 8 (11,1-24) | 31.9 (38.8,4.5-130.2) |
| Cohort 5 | 12 | 63 (14,44-81) | 55 (21,22-65) | 1.7 (6.2,0.1-70.2) | 16 | 67 (18,50-81) | 40 (32,8-65) | 5.2 (66.6,0.3-237.7) |
| Cohort 6 | 3 | 63 (10,52-72) | 65 (0,65-65) | 5 (4.3,0.5-9.1) | 7 | 59 (14,49-66) | 64 (9,45-65) | 1.3 (0.5,0.6-2.2) |
| Cohort 7 | 5 | 61 (4,57-65) | 49 (43,14-63) | 4.3 (1.3,0.3-6.9) | 9 | 53 (10,45-69) | 63 (4,6-65) | 1.5 (1.2,0.04-5.3) |
| Cohort 8 | 4 | 58 (16,45-74) | 50 (10,32-55) | 45.2 (43,0.7-64.6) | 7 | 56 (9,45-68) | 45 (15,25-55) | 35.8 (47.2,0.9-103.9) |
| Cohort 9 | 3 | 59 (1,58-60) | 42 (11,26-49) | 2.5 (11.6,0.8-24) | 8 | 59 (5,45-68) | 36 (20,15-49) | 4.9 (22,0.7-53.7) |
| Cohort 10 | 4 | 61 (11,46-73) | 55 (9,35-58) | 10.9 (28,0.4-59) | 4 | 57 (4,53-64) | 43 (5,37-59) | 39.2 (16.9,1.4-55.9) |
| Cohort 11 | 6 | 65 (5,51-75) | 47 (24,23-61) | 5.2 (31.3,0.4-201.4) | 16 | 59 (10,23-75) | 49 (18,29-64) | 5.8 (42,0.6-71) |
| Cohort 12 | 4 | 51 (16,32-62) | 58 (8,54-63) | 1.8 (14.5,0.3-55.9) | 8 | 58 (9,47-80) | 51 (11,43-62) | 6.5 (9,1.3-34.3) |
| Cohort 13 | 4 | 68 (20,31-75) | 60 (4,51-62) | 23.2 (55,0.5-107.5) | 8 | 66 (16,52-83) | 40 (27,19-61) | 3.4 (27,0.2-62.6) |
| Cohort 14 | 18 | 49 (14,36-79) | 41 (15,24-47) | 14.3 (45.9,0.7-143.6) | 10 | 54 (15,25-76) | 43 (9,31-53) | 3.1 (10.8,0.1-16.5) |
| Cohort 15 | 3 | 47 (11,42-63) | 18 (5,11-21) | 36.7 (91.4,3.2-186.1) | 7 | 62 (12,51-76) | 35 (16,23-60) | 8.6 (3.3,3-97.5) |
| Cohort 16 | 14 | 68 (12,43-93) | 48 (33,20-65) | 12.1 (24.9,1.1-54.4) | 26 | 65 (9,45-81) | 47 (26,4-62) | 8 (25.2,0.5-111.8) |
| Cohort 17 | 15 | 68 (19,37-79) | 38 (34,8-64) | 7.6 (31.2,0.5-188.4) | 20 | 72 (12,51-80) | 56 (11,23-65) | 5.2 (22,0.3-110.8) |
| Cohort 18 | 12 | 62 (15,34-85) | 28 (5,14-34) | 15.6 (47.9,0.8-136.9) | 15 | 64 (14,50-78) | 27 (4,19-34) | 4.8 (18.8,0.7-100.4) |
| **Total** | **134** | **61 (20,30-93)** | **39 (29,0-65)** | **9.7(42.1,0.1-201.4)** | **215** | **62 (16,23-83)** | **43 (26,1-65)** | **5.8 (26.9,0.04-237.7)** |

**Supplemental Table 3.** Summary statistics from robust mixed-effects linear regression to test associations between ipsilesional hippocampal volume and sensorimotor impairment (*top*) and contralesional hippocampal volume and sensorimotor impairment (*bottom*) in participants who passed quality control for bilateral hippocampi. The full model as well as the sample size (*N*), conditional $R^2$, beta coefficient (*Beta*) with 95% confidence interval (*CI*), standard error (*SE*), *t-value* and degrees of freedom *t*(*DF*), standardized *d-value*, uncorrected *p-value* for all fixed effect covariates are reported. Significant covariates are denoted in bold.

| ***Hippocampus ~ FMA-UE + Sex + Lesioned Hemisphere + Age + random(Cohort)*** | | | | | |
|---|---|---|---|---|---|
| ***Covariates*** | ***Beta(CI)*** | ***SE*** | ***t(DF)*** | ***d-value*** | ***p-value*** |
| ***IPSILESIONAL HIPPOCAMPAL VOLUME (N=334; $R^2$=0.27)*** | | | | | |
| **FMA-UE** | 0.16 (0.05 – 0.27) | 0.06 | 2.86(285) | 0.34 | **0.004** |
| **Sex** | -0.54 (-0.74 – -0.34) | 0.10 | -5.29(323) | -0.59 | **<0.001** |
| Lesioned Hemisphere | 0.19 (-0.01 – 0.39) | 0.10 | 1.86(334) | 0.20 | 0.06 |
| **Age** | -0.32 (-0.43 – -0.22) | 0.05 | -6.16(334) | -0.67 | **<0.001** |
| ***CONTRALESIONAL HIPPOCAMPAL VOLUME (N=334; $R^2$=0.33)*** | | | | | |
| FMA-UE | 0.02 (-0.09 – 0.12) | 0.05 | 2.86(285) | 0.34 | 0.78 |
| **Sex** | -0.55 (-0.75 – -0.36) | 0.10 | -5.29(323) | -0.59 | **<0.001** |
| **Lesioned Hemisphere** | -0.36 (-0.56 – -0.16) | 0.10 | 1.86(334) | 0.20 | **0.001** |
| **Age** | -0.41 (-0.51 – 0.32) | 0.05 | -6.16(334) | -0.67 | **<0.001** |

**Supplemental Table 4.** Summary statistics from robust mixed-effects linear regression to test associations between ipsilesional hippocampal volume and sensorimotor impairment (*top*) and contralesional hippocampal volume and sensorimotor impairment (*bottom*) when including a sensorimotor impairment and sex interaction in participants who passed quality control for bilateral hippocampi. The full model as well as the sample size (*N*), conditional $R^2$, beta coefficient (*Beta*) with 95% confidence interval (*CI*), standard error (*SE*), *t-value* and degrees of freedom *t*(*DF*), standardized *d-value*, uncorrected *p-value* for all fixed effect covariates are reported. Significant covariates are denoted in bold.

| ***Hippocampus ~ FMA-UE*Sex + FMA-UE + Sex + Lesioned Hemisphere + Age + random(Cohort)*** | | | | | |
|---|---|---|---|---|---|
| ***Covariates*** | ***Beta(CI)*** | ***SE*** | ***t(DF)*** | ***d-value*** | ***p-value*** |
| ***IPSILESIONAL HIPPOCAMPAL VOLUME (N=334; $R^2$=0.29)*** | | | | | |
| **FMA-UE** | 0.32 (0.16 – 0.47) | 0.08 | 3.97(330) | 0.44 | **<0.001** |
| **FMA-UE*Sex** | -0.27 (-0.47 – -0.07) | 0.10 | -2.68(330) | -0.30 | **0.007** |
| **Sex** | -0.54 (-0.74 – -0.35) | 0.10 | -5.36(323) | -0.60 | **<0.001** |
| Lesioned Hemisphere | 0.18 (-0.03 – 0.38) | 0.10 | 1.71(334) | 0.19 | 0.09 |
| **Age** | -0.32 (-0.43 – -0.22) | 0.05 | -6.24(334) | -0.68 | **<0.001** |
| ***CONTRALESIONAL HIPPOCAMPAL VOLUME (N=334; $R^2$=0.35)*** | | | | | |
| FMA-UE | 0.17 (0.02 – 0.32) | 0.08 | 2.20 (322) | 0.25 | 0.028 |
| **FMA-UE*Sex** | -0.27 (-0.46 – -0.08) | 0.10 | -2.77(333) | -0.30 | **0.006** |
| **Sex** | -0.56 (-0.76 – -0.37) | 0.10 | -5.73(327) | -0.63 | **<0.001** |
| **Lesioned Hemisphere** | -0.38 (-0.58 – -0.19) | 0.10 | -3.85(332) | -0.42 | **<0.001** |
| **Age** | -0.42 (-0.51 – -0.32) | 0.05 | -8.29(332) | -0.91 | **<0.001** |

**Supplemental Table 5.** Summary statistics from robust mixed-effects linear regression to test associations between ipsilesional hippocampal volume and sensorimotor impairment (*top*) and contralesional hippocampal volume and sensorimotor impairment (*bottom*) when including lesion size as a covariate in participants who passed quality control for bilateral hippocampi. The full model as well as the sample size (*N*), conditional $R^2$, beta coefficient (*Beta*) with 95% confidence interval (*CI*), standard error (*SE*), *t-value* and degrees of freedom *t*(*DF*), standardized *d-value*, uncorrected *p-value* for all fixed effect covariates are reported. Significant covariates are denoted in bold.

| ***Hippocampus ~ Lesion Size + FMA-UE*Sex + FMA-UE + Sex + Lesioned Hemisphere + Age + random(Cohort)*** | | | | | |
|---|---|---|---|---|---|
| ***Covariates*** | ***Beta(CI)*** | ***SE*** | ***t(DF)*** | ***d-value*** | ***p-value*** |
| ***IPSILESIONAL HIPPOCAMPAL VOLUME (N=334; $R^2$=0.32)*** | | | | | |
| **FMA-UE** | 0.27 (0.11 – 0.42) | 0.08 | 3.38(330) | 0.37 | **0.001** |
| **FMA-UE*Sex** | -0.27 (-0.46 – -0.08) | 0.10 | -2.74(330) | -0.30 | **0.006** |
| **Lesion Size** | -0.19 (-0.30 – -0.09) | 0.05 | -3.75(331) | -0.41 | **<0.001** |
| **Sex** | -0.60 (-0.79 – -0.40) | 0.10 | -5.97(323) | -0.66 | **<0.001** |
| Lesioned Hemisphere | 0.17 (-0.03 – 0.37) | 0.10 | 1.71(334) | 0.19 | 0.09 |
| **Age** | -0.36 (-0.46 – -0.26) | 0.05 | -7.01(334) | -0.77 | **<0.001** |
| ***CONTRALESIONAL HIPPOCAMPAL VOLUME (N=334; $R^2$=0.34)*** | | | | | |
| FMA-UE | 0.17 (0.01 – 0.32) | 0.08 | 2.15(323) | 0.24 | 0.032 |
| **FMA-UE*Sex** | -0.27 (-0.46 – -0.08) | 0.10 | -2.77(333) | -0.30 | **0.006** |
| Lesion Size | -0.01 (-0.11 – 0.09) | 0.05 | -0.20(333) | -0.02 | 0.84 |
| **Sex** | -0.57 (-0.76 - -0.37) | 0.10 | -5.69(327) | -0.63 | **<0.001** |
| **Lesioned Hemisphere** | -0.38 (-0.52 – -0.32) | 0.10 | -3.85(332) | -0.42 | **<0.001** |
| **Age** | -0.42 (-0.52 – -0.32) | 0.05 | -8.21(332) | -0.90 | **<0.001** |

**Supplemental Table 6.** Summary statistics from robust mixed-effects linear regression to test associations between ipsilesional hippocampal volume and lesion size (*top*) and contralesional hippocampal volume and lesion size (*bottom*) in participants who passed quality control for bilateral hippocampi. The full model as well as the sample size (*N*), conditional $R^2$, beta coefficient (*Beta*) with 95% confidence interval (*CI*), standard error (*SE*), *t-value* and degrees of freedom *t*(*DF*), standardized *d-value*, uncorrected *p-value* for all fixed effect covariates are reported. Significant covariates are denoted in bold.

| ***Hippocampus ~ Lesion Size + Sex + Lesioned Hemisphere + Age + random(Cohort)*** | | | | | |
|---|---|---|---|---|---|
| ***Covariates*** | ***Beta(CI)*** | ***SE*** | ***t(DF)*** | ***d-value*** | ***p-value*** |
| ***IPSILESIONAL HIPPOCAMPAL VOLUME (N=334; $R^2$=0.32)*** | | | | | |
| **Lesion Size** | -0.22 (-0.32 – -0.12) | 0.05 | -4.24(333) | -0.46 | **<0.001** |
| **Sex** | -0.59 (-0.79 – -0.39) | 0.10 | -5.84(322) | -0.65 | **<0.001** |
| Lesioned Hemisphere | 0.17 (-0.03 – 0.36) | 0.10 | 1.66(334) | 0.18 | 0.10 |
| **Age** | -0.35 (-0.45 – -0.25) | 0.05 | -6.79(334) | -0.74 | **<0.001** |
| ***CONTRALESIONAL HIPPOCAMPAL VOLUME (N=334; $R^2$=0.33)*** | | | | | |
| Lesion Size | -0.01 (-0.11 – 0.09) | 0.05 | -0.24(334) | -0.03 | 0.81 |
| **Sex** | -0.56 (-0.75 – -0.36) | 0.10 | -5.53(327) | -0.61 | **<0.001** |
| **Lesioned Hemisphere** | -0.36 (-0.56 – -0.17) | 0.10 | -3.65(333) | -0.40 | **0.001** |
| **Age** | -0.42 (-0.52 – -0.32) | 0.05 | -8.10(332) | -0.89 | **<0.001** |

**Supplemental Table 7.** Summary statistics from robust mixed-effects linear regression to test associations between ipsilesional hippocampal volume and sensorimotor impairment (*top*) and contralesional hippocampal volume and sensorimotor impairment (*bottom*) when including lesion size as a covariate and excluding participants with secondary lesions. The full model as well as the sample size (*N*), conditional $R^2$, beta coefficient (*Beta*) with 95% confidence interval (*CI*), standard error (*SE*), *t-value* and degrees of freedom *t*(*DF*), standardized *d-value*, uncorrected *p-value* for all fixed effect covariates are reported. Significant covariates are denoted in bold.

| ***Hippocampus ~ Lesion Size + FMA-UE*Sex + Lesioned Hemisphere + Age + random(Cohort)*** | | | | | |
|---|---|---|---|---|---|
| ***Covariates*** | ***Beta(CI)*** | ***SE*** | ***t(DF)*** | ***d-value*** | ***p-value*** |
| ***IPSILESIONAL HIPPOCAMPAL VOLUME (N=240; $R^2$=0.35)*** | | | | | |
| **FMA-UE** | 0.30 (0.12 – 0.48) | 0.09 | 3.27(239) | 0.42 | **0.001** |
| **FMA-UE*Sex** | -0.31 (-0.53 – -0.08) | 0.11 | -2.67(236) | -0.35 | **0.008** |
| **Lesion Size** | -0.14 (-0.27 – -0.02) | 0.06 | -2.34(237) | -0.30 | **0.019** |
| **Sex** | -0.57 (-0.80 – -0.35) | 0.11 | -5.03(231) | -0.66 | **<0.001** |
| Lesioned Hemisphere | 0.18 (-0.06 – 0.41) | 0.12 | 1.49(240) | 0.19 | 0.14 |
| **Age** | -0.29 (-0.41 – -0.18) | 0.06 | -4.90(237) | -0.64 | **<0.001** |
| ***CONTRALESIONAL HIPPOCAMPAL VOLUME (N=245 $R^2$=0.32)*** | | | | | |
| FMA-UE | 0.18 (0.00 – 0.35) | 0.09 | 1.93(240) | 0.25 | 0.05 |
| **FMA-UE*Sex** | -0.28 (-0.50 – -0.05) | 0.12 | -2.38(243) | -0.31 | **0.017** |
| Lesion Size | 0.05 (-0.08 – 0.17) | 0.06 | 0.75(244) | 0.10 | 0.45 |
| **Sex** | -0.45 (-0.68 – -0.22) | 0.12 | -3.90(240) | -0.50 | **<0.001** |
| **Lesioned Hemisphere** | -0.38 (-0.62 – -0.15) | 0.12 | -3.21 (242) | -0.41 | **0.001** |
| **Age** | -0.36 (-0.48 – -0.25) | 0.06 | -6.05(245) | -0.77 | **<0.001** |

**Supplemental Table 8.** Summary statistics from robust mixed-effects linear regression to test associations between ipsilesional hippocampal volume and lesion size (*top*) and contralesional hippocampal volume and lesion size (*bottom*) after excluding participants with secondary lesions. The full model as well as the sample size (*N*), conditional $R^2$, beta coefficient (*Beta*) with 95% confidence interval (*CI*), standard error (*SE*), *t-value* and degrees of freedom *t*(*DF*), standardized *d-value*, uncorrected *p-value* for all fixed effect covariates are reported. Significant covariates are denoted in bold.

| ***Hippocampus ~ Lesion Size + Sex + Lesioned Hemisphere + Age + random(Cohort)*** | | | | | |
|---|---|---|---|---|---|
| ***Covariates*** | ***Beta(CI)*** | ***SE*** | ***t(DF)*** | ***d-value*** | ***p-value*** |
| ***IPSILESIONAL HIPPOCAMPAL VOLUME (N=240; $R^2$=0.33)*** | | | | | |
| **Lesion Size** | -0.18 (-0.30 – -0.06) | 0.06 | -2.94(239) | -0.38 | **0.003** |
| **Sex** | -0.58 (-0.80 – -0.35) | 0.12 | -5.01(230) | -0.66 | **<0.001** |
| Lesioned Hemisphere | 0.16 (-0.08 – 0.39) | 0.12 | 1.30(240) | 0.17 | 0.19 |
| **Age** | -0.28 (-0.40 – -0.16) | 0.06 | -4.65(237) | -0.60 | **<0.001** |
| ***CONTRALESIONAL HIPPOCAMPAL VOLUME (N=245; $R^2$=0.29)*** | | | | | |
| Lesion Size | 0.04 (-0.08 – 0.16) | 0.06 | 0.62(243) | 0.08 | 0.53 |
| **Sex** | -0.45 (-0.68 – -0.22) | 0.12 | -3.84(240) | -0.50 | **<0.001** |
| **Lesioned Hemisphere** | -0.37 (-0.60 – -0.14) | 0.12 | -3.10(242) | -0.40 | **0.002** |
| **Age** | -0.36 (-0.48 – -0.24) | 0.06 | -5.91(245) | -0.76 | **<0.001** |

#### *Supplementary Figures*

**Supplemental Figure 1.** Lesion density maps for lesions from participants with cohort-reported left and right hemisphere lesions, and without any secondary lesions (e.g., bilateral, brainstem or cerebellar lesions), overlaid on the MNI-152 template.

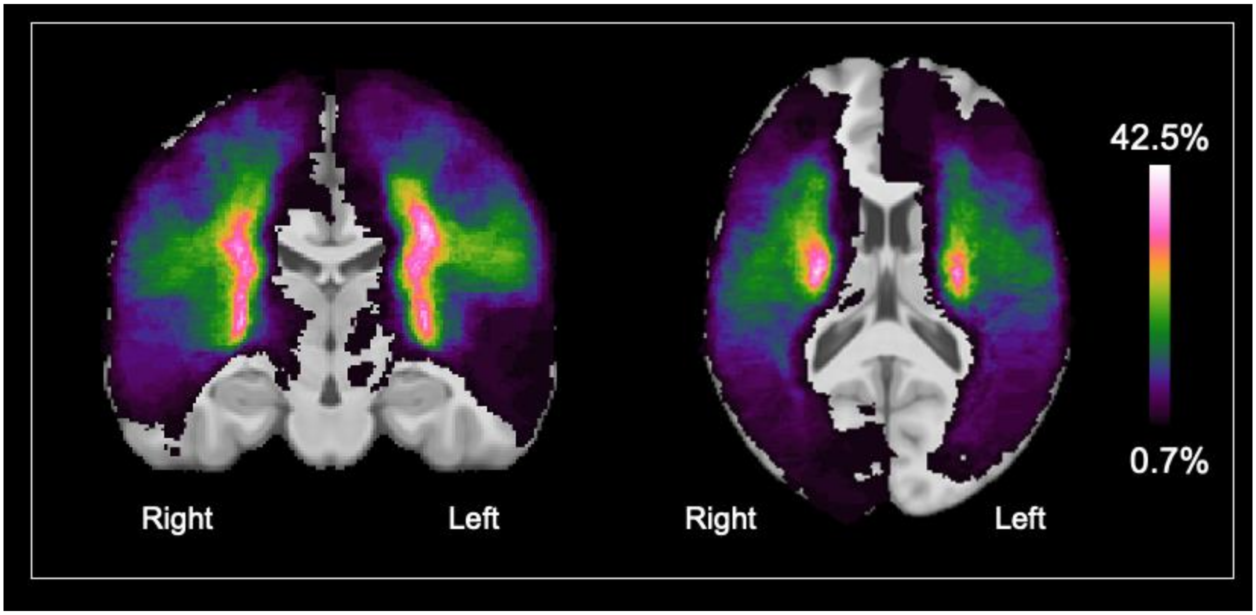
